# Single-molecule diffusivity quantification in *Xenopus* egg extracts elucidates physicochemical properties of the cytoplasm

**DOI:** 10.1101/2024.08.24.609541

**Authors:** Alexander A. Choi, Coral Y. Zhou, Ayana Tabo, Rebecca Heald, Ke Xu

## Abstract

The living cell creates a unique internal molecular environment that is challenging to characterize. By combining single-molecule displacement/diffusivity mapping (SM*d*M) with physiologically active extracts prepared from *Xenopus laevis* eggs, we sought to elucidate molecular properties of the cytoplasm. Quantification of the diffusion coefficients of 15 diverse proteins in extract showed that, compared to in water, negatively charged proteins diffused ∼50% slower, while diffusion of positively charged proteins was reduced by ∼80-90%. Adding increasing concentrations of salt progressively alleviated the suppressed diffusion observed for positively charged proteins, signifying electrostatic interactions within a predominately negatively charged macromolecular environment. To investigate the contribution of RNA, an abundant, negatively charged component of cytoplasm, extracts were treated with ribonuclease, which resulted in low diffusivity domains indicative of aggregation, likely due to the liberation of positively charged RNA-binding proteins such as ribosomal proteins, since this effect could be mimicked by adding positively charged polypeptides. Interestingly, negatively charged proteins of different sizes showed similar diffusivity suppression in extract, which are typically prepared under conditions that inhibit actin polymerization. Restoring or enhancing actin polymerization progressively suppressed the diffusion of larger proteins, recapitulating behaviors observed in cells. Together, these results indicate that molecular interactions in the crowded cell are defined by an overwhelmingly negatively charged macromolecular environment containing cytoskeletal networks.

**Significance Statement:** The complex intracellular molecular environment is notably challenging to elucidate and recapitulate. *Xenopus* egg extracts provide a native yet manipulatable cytoplasm model. Through single-molecule microscopy, here we decipher the cytoplasmic environment and molecular interactions by examining the diffusion patterns of diverse proteins in *Xenopus* egg extracts with strategic manipulations. These experiments reveal an overwhelmingly negatively charged macromolecular environment with crosslinked meshworks, offering new insight into the inner workings of the cell.

## Introduction

The living cell creates a unique molecular environment within its micrometer-sized bound volume (*1-4*). This milieu is notably difficult to recapitulate *in vitro*: Simple buffers, including those added with inert macromolecules to emulate the crowded intracellular environment, often do not maintain the native protein interactions or achieve the optimal activities seen in the cell. A precise mix of proteins, nucleic acids, peptides, as well as small molecules and ions at the correct concentrations appears necessary to attain the magic of life. How this myriad of molecular species interacts with each other remains a mystery.

Diffusion measurements provide a valuable window into molecular environments and interactions (*5-7*). With the development of single-molecule displacement/diffusivity mapping (SM*d*M), a high-precision diffusion quantification method based on high-throughput single-molecule microscopy and statistics (*8, 9*), we have recently reported that for fluorescent proteins (FPs) expressed in mammalian cells, the translational diffusion of positively, but not negatively, charged species is substantially impeded in diverse cytoplasmic and intraorganellar environments (*8, 10*). Together with other recent microscopy and NMR studies noting reduced diffusivities of positively charged proteins in bacterial and mammalian cells (*11-15*), these results point to an intriguing, charge sign-asymmetric intracellular environment.

Expression of FPs in the cell, however, offers limited control over the types and amounts of diffusers that can be examined, and nuances in diffusivity may be masked by heterogeneity due to intracellular structures and cell-to-cell variations. We have thus also examined protein interactions in solution, and demonstrated that simple negatively charged proteins like bovine serum albumin (BSA) biasedly impede the diffusion of positively charged proteins in an ionic strength-dependent fashion (*16*). Although easy to manipulate, in-solution experiments are limited by their simplicity and artificial nature.

*Xenopus laevis* egg extracts present an opportunity to bridge the above two limits. Prepared as homogenized, undiluted cytoplasm, egg extracts are an excellent model that retains the native intracellular environment while being highly manipulatable through simple biochemical approaches (*17-19*). Many complex cellular processes have been recapitulated and/or studied in this system, including cell cycle timing, mitotic spindle formation, chromosome condensation, cell-like compartmentalization, and apoptosis (*18-26*). Several recent studies have highlighted the use of *Xenopus* egg extracts to interrogate the biophysical properties of the cytoplasm (*27-29*), in which molecular diffusion plays key roles.

Capitalizing on the versatility of *Xenopus laevis* egg extracts and the high precision of SM*d*M, here we quantify the diffusion behaviors of diverse proteins both in the native cytoplasmic environment and after strategic manipulations. By examining proteins spanning a wide range of sizes and charges and modulating the extract ionic strength, we first show that the egg cytoplasm is an overwhelmingly negatively charged macromolecular environment. Through ribonuclease treatments, we next uncover the role of RNA in neutralizing positively charged ribosomal proteins to prevent cytoplasmic aggregation. Finally, by inhibiting or enhancing actin polymerization, we underscore its key role in the size-dependent regulation of diffusion in the cytoplasm.

## Results and Discussion

*Xenopus laevis* egg cytoplasmic extracts (hereafter “extracts”) were prepared following standard protocols (**Fig. 1a** and Methods) (*17*). Dye-labeled proteins (Methods and **Table S1**) were added to the extract at <1 nM, optimal for SM*d*M single-molecule detection and diffusion quantification while minimizing possible interactions between the added molecules (*16*). For SM*d*M (Methods) (*8, 30*), the sample was illuminated with an excitation laser at a depth of ∼3 μm for the wide-field recording of single-molecule images with an EM-CCD camera. The laser was repeatedly modulated as paired stroboscopic pulses across tandem camera frames at a fixed center-to-center separation of Δ*t* = 1 ms (**Fig. 1b**), so that transient displacements in the Δ*t* time window were detected for molecules that diffused into the field of view (**Fig. 1c**). The execution of ∼10^4^ frame pairs over ∼3 min thus recorded ∼10^5^ transient single-molecule displacements. The accumulated displacements were either spatially binned for local statistics (*8, 31*) to generate a diffusivity map (**Fig. S1**), or pooled for global fitting to yield a diffusion coefficient *D* with an ∼±1% bracket at 95% confidence (**Fig. 1d,e**) (*30*).

**Figure 1.**
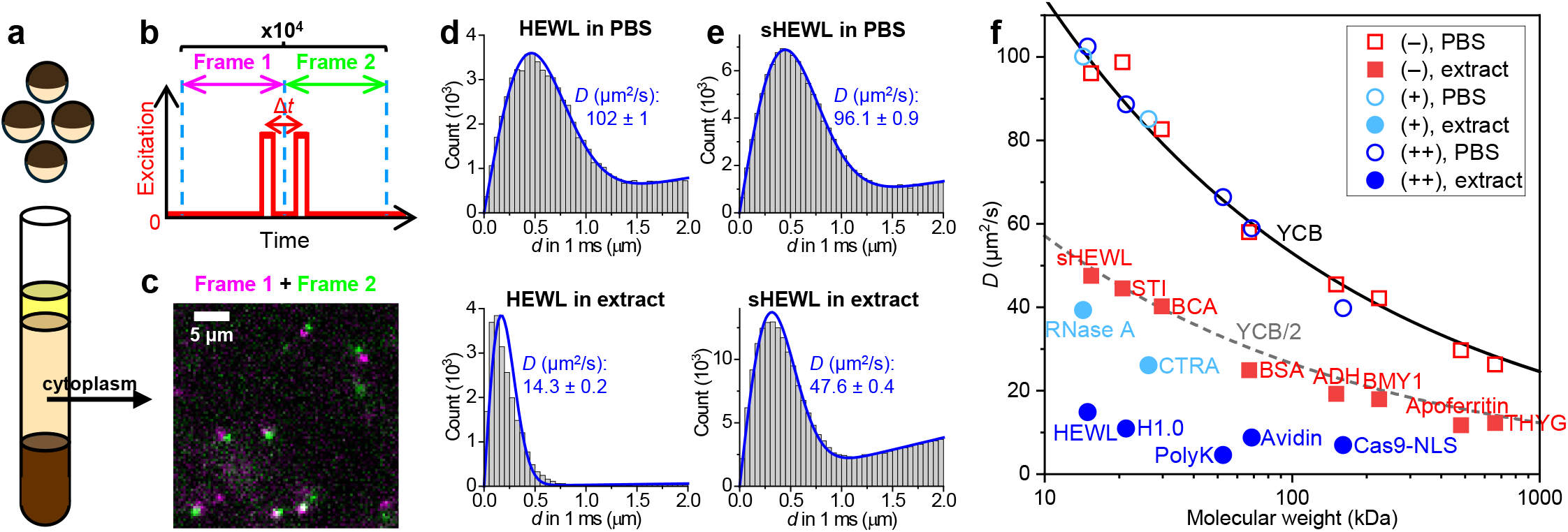
Single-molecule displacement/diffusivity mapping (SM*d*M)-based protein diffusivity quantification in *Xenopus* egg extract versus in phosphate-buffered saline (PBS). (**a**) Schematic: Cytoplasmic extracts from *Xenopus* eggs. (**b**) Schematic: In SM*d*M, paired excitation pulses are repeatedly applied across tandem camera frames, so that transient single-molecule displacements are captured in the wide field for the time window defined by the separation between the paired pulses, Δ*t*. (**c**) Example single-molecule images of Cy3B-labeled bovine carbonic anhydrase diffusing in the extract, shown as a magenta-green overlaid image for a tandem pair of frames at Δ*t* = 1 ms. (**d**) Example distributions of SM*d*M-recorded 1-ms single-molecule displacements for ∼200 pM Cy3B-labeled hen egg white lysozyme (HEWL) diffusing in PBS (top) versus in extract (bottom). Blue curves: fits to our single-mode diffusion model, with resultant apparent diffusion coefficients *D* and 95% confidence intervals marked in each plot. (**e**) Similar to (d), but for succinylated HEWL (sHEWL). (**f**) SM*d*M-determined *D* values for 15 proteins of varied sizes and charges (**Table S1**), in PBS (hollow symbols) and extract (filled symbols). Red squares: Negatively charged proteins. Blue circles: Positively charged proteins with >+5 net charges. Light-blue circles: Weakly positively charged proteins with ∼+2 net charges. Each data point is an average of at least three SM*d*M measurements from two or more extract samples. Solid curve: Expected *D* in PBS at room temperature according to the Young−Carroad−Bell (YCB) model. Dashed curve: The PBS YCB values divided by 2.

**Figure 1d** compares SM*d*M results of Cy3B-tagged hen egg white lysozyme (HEWL) in phosphate-buffered saline (PBS) versus in extract. With a net charge of ∼+7 at physiological pH, the ∼15 kDa protein serves as a model to examine charge interactions (*16*). In PBS, the SM*d*M-recorded single-molecule displacements fit well to our single-mode diffusion model to yield a diffusion coefficient *D* = 102 μm^2^/s (**Fig. 1d**), consistent with previous results (*16, 30*). In contrast, markedly suppressed displacements were observed in the extract, which conformed less well to the single-mode model to yield a substantially lower apparent *D* value of 14 μm^2^/s (**Fig. 1d**). Spatial mapping of *D* showed no noticeable features (**Fig. S1a**), expected for the homogenized extract, and suggests the above-observed deviation from the single-mode diffusion model was due not to spatial variations but likely to heterogeneous transient interactions with extract components. Fitting the SM*d*M displacements to two modes (**Fig. S2**) yielded slow and fast components as 3.4 and 21.3 μm^2^/s, respectively, although a continuous distribution of different transient states is likely. To understand if the positive net charge of HEWL drove the above behavior, we succinylated HEWL to shift its net charge from ∼+7 to ∼−4 (Methods and **Table S1**) (*16, 32*). SM*d*M yielded a notably higher *D* of 48 μm^2^/s for succinylated HEWL (sHEWL) in the extract with a good fit to the single-mode diffusion model (**Fig. 1e**).

We next quantified and compared the diffusivity of 15 soluble proteins of diverse sizes and net charges (**Table S1**), many of which have been employed as protein size standards, in PBS versus extract. As we plotted all *D* values in PBS (hollow symbols in **Fig. 1f**) against the protein molecular weight, we noted a monotonic decrease in agreement with the Young-Carroad-Bell (YCB) model (solid curve in **Fig. 1f**) (*33*). This result is expected: The YCB model is based on fitting experimental diffusivity values, and common proteins have exhibited *D* values within 10% of the YCB prediction (*33, 34*). Our previous SM*d*M results also well followed the YCB trend (*30, 35*).

For diffusion in the extract, the SM*d*M-determined *D* values of the 8 negatively charged soluble proteins (filled red squares in **Fig. 1f**) showed a size dependence that roughly followed (or were slightly lower than) 50% of the YCB values in PBS (dashed curve in **Fig. 1f**). An early NMR study reports a uniform 45% scaling of the apparent diffusion coefficients of small molecules in *Xenopus* oocytes versus in water (*36*). Using a Cannon-Fenske viscometer, we measured the bulk viscosity of the extract as 2.30 cP, ∼2.22x of that measured for PBS (1.03 cP). Thus, the diffusion of negatively charged proteins is only passively impeded by the higher cytoplasm viscosity over water. In comparison, the 7 positively charged soluble proteins (filled blue and light-blue circles in **Fig. 1f**) exhibited further suppressions in diffusivity well below the ∼50% PBS values.

Plotting the SM*d*M-determined *D* values in the extract relative to those in PBS as a function of the protein net charge (**Fig. 2a**) showed that whereas the negatively charged proteins all diffused at 40-50% of their diffusivities in PBS, proteins carrying >+5 net charges diffused at 10-20% of their PBS diffusivities, while the weakly positively charged proteins exhibited intermediate values.

**Figure 2.**
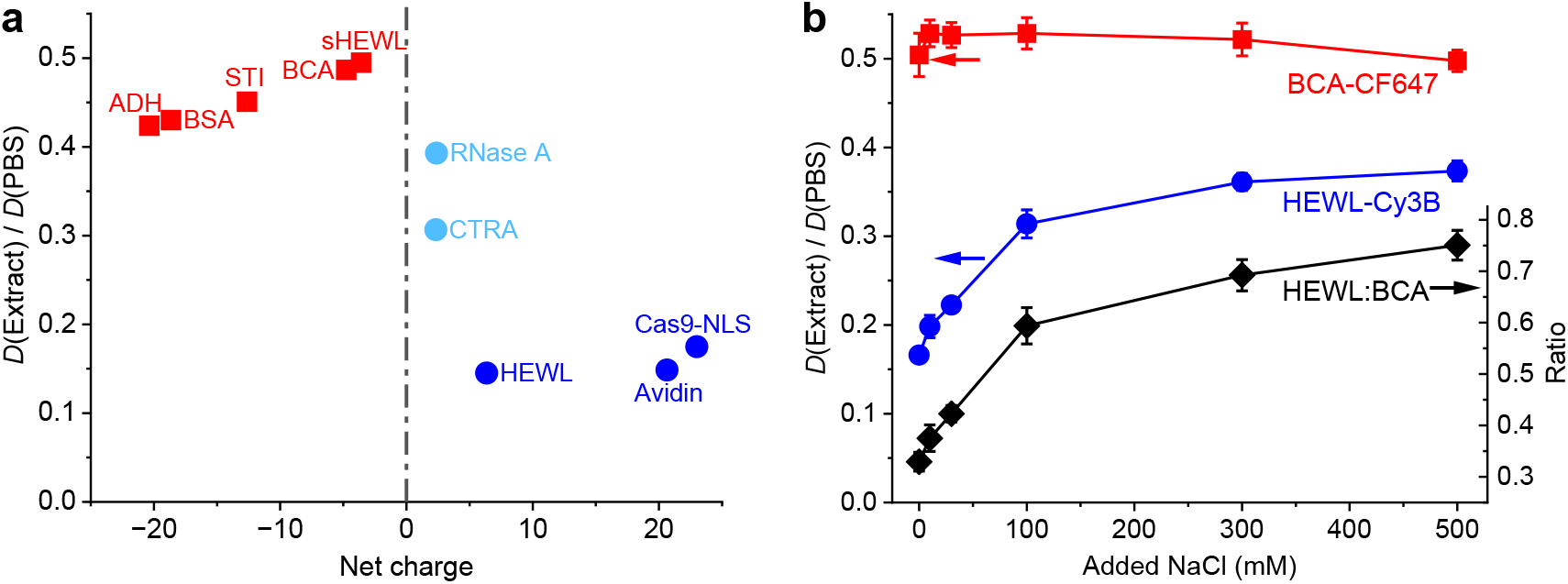
Net-charge effects on protein diffusion in *Xenopus* egg extract. (**a**) SM*d*M-determined diffusion coefficients *D* in the extract relative to those in PBS for different proteins, as a function of their net charges in the range of -25 to +25. (**b**) Blue and red symbols: Relative in-extract *D* values normalized to in-PBS values, plotted as a function of added NaCl, for the positively charged HEWL-Cy3B (blue) and the negatively charged BCA-CF647 (red). Values were obtained through sequential SM*d*M in two color channels. Error bars: Sample standard deviations between results from two or three SM*d*M measurements at each data point. Black diamonds (y-axis on the right): Ratio between the PBS-normalized *D* values of the two proteins.

To verify electrostatic interaction as the driving force of this behavior, we added increasing concentrations of salt to the extract and re-measured *D*. To directly compare positively and negatively charged proteins in the same extract samples, we separately labeled HEWL and bovine carbonic anhydrase (BCA) with Cy3B and CF647, respectively, and performed SM*d*M for both proteins in two color channels. We found that for samples with increasing amounts of NaCl added, the positively charged HEWL (blue circles in **Fig. 2b**) progressively increased its diffusivity from 16% to 37% of the PBS value. In contrast, the negatively charged BCA (red squares in **Fig. 2b**) stayed at 50-53% of its PBS value. Assuming the remaining minor variations in the BCA readouts reflected non-charge effects, *e*.*g*., mechanical disturbances from NaCl addition, the ratio between the PBS-normalized *D* values of HEWL and BCA (black diamonds in **Fig. 2b**) progressively increased with NaCl addition from 10 mM to 300 mM before plateauing at ∼70%. Thus, ionic screening efficiently reduced the likely dynamic interactions between the positively charged HEWL and the negatively charged species in the extract, yet the interactions were still not fully eliminated at 500 mM ionic strength, reminiscent of our recent observations of HEWL-BSA interactions in buffer (*16*).

Together, our SM*d*M experiments in the extract unveiled charge-driven suppression of diffusion for positively charged proteins. These results may be explained if the macromolecular environment in the extract is dominated by negatively charged species, akin to our recent observations in mammalian cells and in solution (*8, 10, 16*). In agreement with this prediction, an analysis of the expected net charges of major proteins in the *Xenopus* egg cytoplasmic extract, as ranked by the mass spectrometry-detected abundances (*37*), indicated most as negative or neutral (**Table S2**). Cytoplasmic extracts also contain abundant ribosomes. Although many ribosomal proteins are highly positively charged (**Table S3**), their assembly with ribosomal RNA (rRNA) results in highly negatively charged ribosomes, which in bacteria have been identified as the source of diffusion suppression for positively charged proteins (*11*).

To probe the contribution of RNA, ∼90% of which is ribosomal in the extract (*38*), to protein diffusion properties, we treated extracts with ribonuclease (RNase) (*29, 39, 40*). Interestingly, we observed that while treating extracts with RNase A did not immediately alter the diffusion behavior of the positively charged HEWL (**Fig. 3a-b**, versus **Fig. S1a-b**), micrometer-scale low-diffusivity domains emerged after 3 h (magenta in **Fig. 3c**). Rendering the single-molecule localizations from the SM*d*M data into SMLM (single-molecule localization microscopy) (*41*) super-resolution images showed an increased presence of HEWL in the low-diffusivity domains and resolved structures consistent with amorphous aggregates with micrometer-scale clouds and nanoscale foci (**Fig. S3**). Meanwhile, the negatively charged BCA also exhibited locally reduced diffusivity in the RNase A-induced aggregates but was moderately excluded (**Fig. S4**). Concomitantly, we noted that the RNase A-treated extracts became visibly cloudy (**Fig. 3e**), consistent with past studies showing that RNase treatments of mammalian and bacterial lysates cause widespread protein aggregation (*42, 43*). Given the high abundance of ribosomal proteins and rRNA in egg extract (*38, 44*), we estimated that RNA degradation may release up to 5 mg/mL ribosomal proteins, many of which are highly positively charged (**Table S3**), into the negatively charged cytoplasmic background. This extensive mixing of oppositely charged proteins likely induces aggregation (*45-47*).

**Figure 3.**
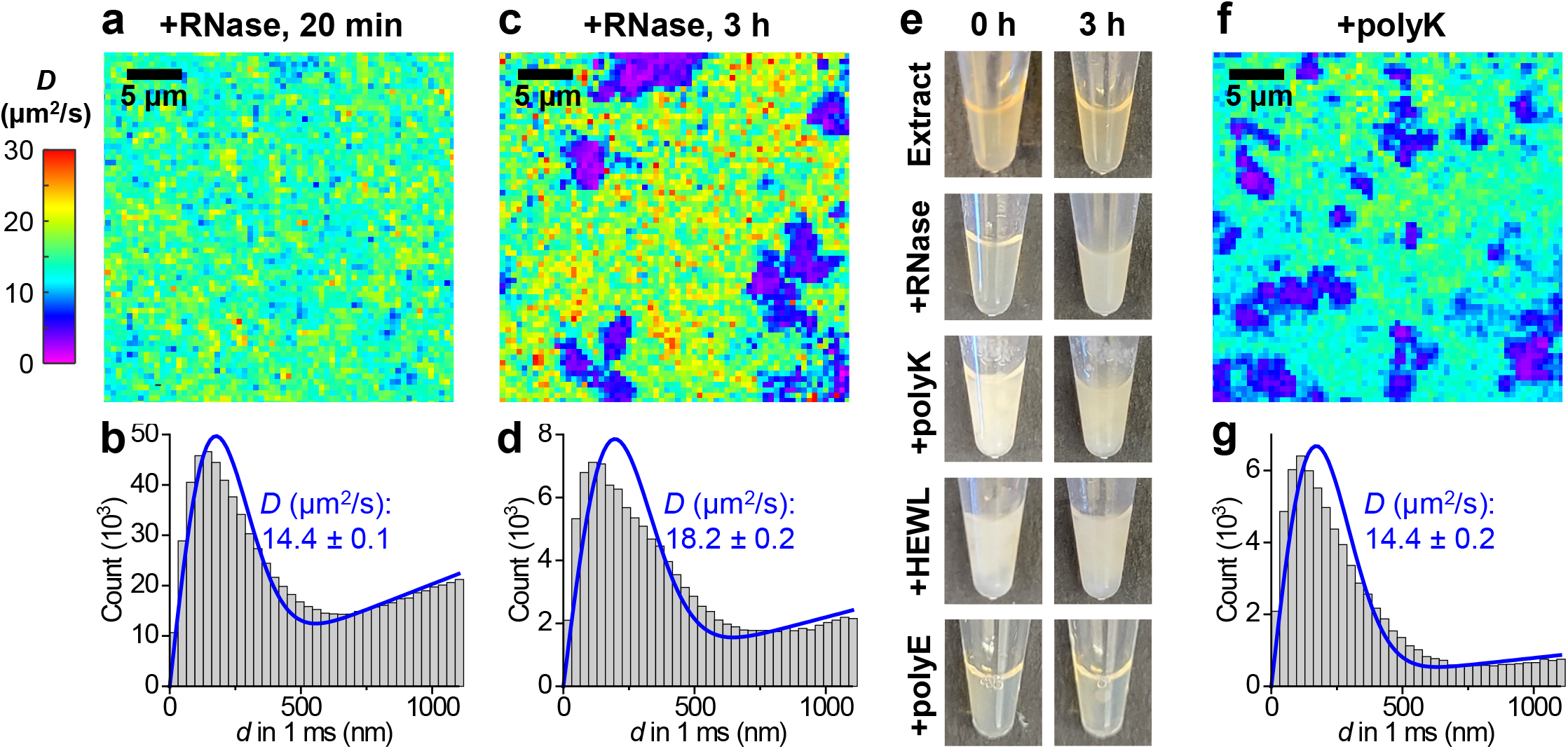
SM*d*M diffusivity mapping of HEWL in RNase-treated egg extracts further underscores the charge-sign asymmetry of the cytoplasmic environment. (**a**) Color-coded SM*d*M diffusivity map of Cy3B-labeled HEWL in an extract sample 20 min after RNase A treatment at room temperature. (**b**) Distributions of single-molecule displacements for the data in (a). Blue line: fit to the SM*d*M diffusion model, with resultant apparent diffusion coefficient *D* and 95% confidence intervals noted. (**c**) SM*d*M diffusivity map of the same sample after 3 h. (**d**) Distributions of single-molecule displacements for regions outside the low-diffusivity domains in (c). (**e**) Photos of extract samples taken immediately (left) and after 3 h (right), after RNase A treatment or the addition of 1 mg/mL polylysine, HEWL, or polyglutamic acid. (**f**) Color-coded SM*d*M *D* map of Cy3B-labeled HEWL in an extract sample supplemented with 1 mg/mL polylysine. (**g**) Distribution of single-molecule displacements for regions outside the low-diffusivity domains in (f).

To examine this hypothesis, we added positively charged proteins to untreated extracts. Notably, addition of 1 mg/mL of either polylysine or HEWL induced immediate clouding of the extract (**Fig. 3e**) and micrometer-sized low-diffusivity domains in the SM*d*M diffusivity map (**Fig. 3f**). In comparison, adding negatively charged polyglutamic acid (**Fig. 3e**) or BSA (**Fig. S1b**) to the extract did not induce aggregation. To further substantiate our model, we examined high-speed extracts (HSEs) in which ribosomes (together with vesicles and large protein complexes) were removed by ultracentrifugation at 200,000*g* for 2.5 h at 4 °C (*17*). RNase treatment of the ribosome-depleted HSE did not induce clouding, while polylysine addition still generated immediate clouding (**Fig. S5**), consistent with our model that the release of ribosomal proteins drove aggregation in the RNase A-treated cytoplasmic extracts.

A recent study showed that introducing cationic “killer peptides” into cell lysates induced aggregation (*48*), echoing earlier findings in bacteria (*49*). Our results suggest that in the cytoplasmic environment, substantial addition or liberation of positively charged proteins, regardless of specific forms, both prompt aggregation. The former scenario is fast due to immediate charge interactions whereas the latter is slow, as positively charged proteins are gradually released. Earlier work showed that RNase treatments of *Xenopus* egg extracts abolish mitotic spindle assembly and nuclear envelope formation (*39, 40*). While these observations, together with the above-noted RNase-induced lysate aggregation (*42, 43*), were interpreted as unrecognized translation-independent RNA functions, our data suggests that a critical RNA function is to invert the positive net charges of their binding proteins to negative, which is essential to maintain a functional cytoplasmic milieu.

In addition to the difference in aggregation speed, the RNase-treated extract also showed a mild increase in *D* of HEWL from ∼14 to ∼18 μm^2^/s for regions outside the low-diffusivity aggregates (compare **Fig. 3d** to **Fig. 3b,g**). This reduced suppression of diffusion may be attributed to diminished impediments from ribosomes and RNA. Yet, the recovery is far from complete, given the above *D* = 48 μm^2^/s of sHEWL in the extract, suggesting that the predominately negatively charged protein environment still suppressed the diffusion of the positively charged HEWL (*16*).

A remaining puzzle was the above-noted relatively invariant 40-50% scaling of *D* in the extract versus in PBS for negatively charged proteins of different sizes (**Fig. 2a**). While this observation simplified our comparison with positively charged diffusers, it conflicts with the notion that intracellular diffusion is generally more hindered for larger molecules (*6, 50-52*). To reconcile this issue, we noted that in the standard extract preparation protocol, cytochalasin is added to inhibit actin polymerization (*17*). It has been shown that the size-dependent suppression of DNA mobility in mammalian cells depends on the actin cytoskeleton, and that in solution, the addition of 8 mg/mL polymerized actin recapitulated a size-dependent diffusion slowdown, whereas adding soluble crowding agents, including cytosol extracts, did not (*50*). Our recent SM*d*M results with expandable hydrogels also showed that obstruction due to immobile meshworks is vital for the size-dependent suppression of molecular diffusivity (*53*). Meanwhile, a recent study reported that BSA diffuses slightly slower in actin-intact extracts (*27*).

To probe the likely role of the actin cytoskeleton in size-dependent diffusion suppression, we prepared actin-intact *Xenopus* egg extracts in which cytochalasin was omitted, as well as actin-supplemented samples to which extra actin was added at 5 mg/mL (∼0.5 wt%), a concentration at which size-dependent diffusion suppression emerges in solution-hydrogel systems (*50, 53*). Chemically fixing the samples for phalloidin labeling (*54*) and three-dimensional stochastic optical reconstruction microscopy (3D-STORM) (*55, 56*) super-resolution imaging revealed dense actin networks with sub-micrometer grid sizes (**Fig. 4a**) analogous to those seen in animal cells (*54*), with the actin-supplemented samples roughly doubling the density of actin filaments over the actin-intact samples. In comparison, no phalloidin staining was observed in the standard actin-inhibited samples (**Fig. S6**).

**Figure 4.**
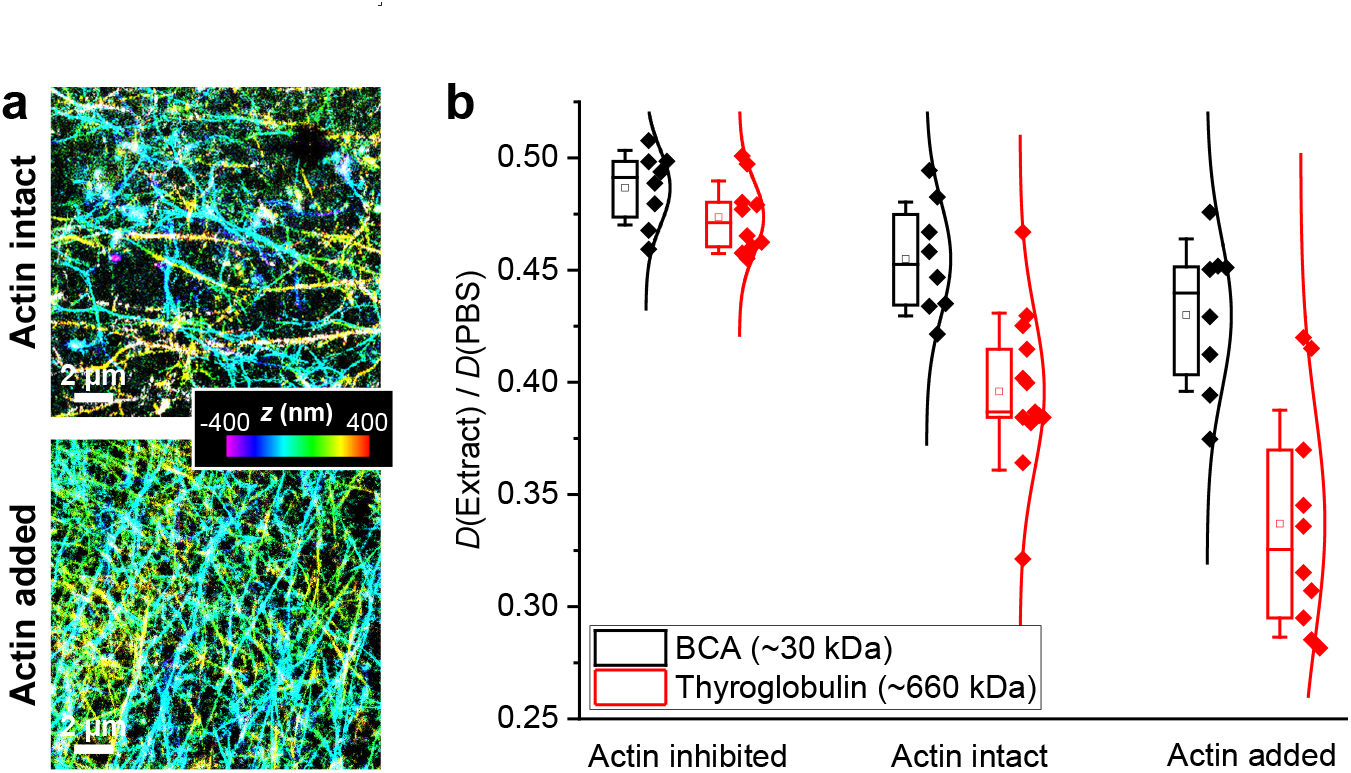
Importance of the actin cytoskeletal network in molecular size-dependent diffusion suppression. (**a**) 3D-STORM super-resolution images of phalloidin-labeled actin filaments in actin-preserved (top) and actin-supplemented (bottom) *Xenopus* egg extracts. Color presents axial (depth) information. (**b**) SM*d*M-determined *D* values in the extract relative to in PBS, for the ∼30 kDa BCA (black) and the ∼660 kDa thyroglobulin (red), in actin-inhibited, actin-intact, and actin-supplemented samples. Each data point corresponds to one independent SM*d*M measurement for a different sample region, from ∼3 samples under each condition.

SM*d*M showed that whereas the 660 kDa thyroglobulin exhibited similar scaling of its *D* value in actin-inhibited extract relative to PBS when compared to the 30 kDa BCA (47% vs. 50%), it experienced progressively stronger diffusivity suppression in actin-intact and actin-supplemented extracts, reaching 39% and 32% of the PBS value, respectively (**Fig. 4b**). In comparison, the 30 kDa BCA displayed notably smaller decreases in *D* to reach 46% and 43% of the PBS value in the actin-intact and actin-supplemented extracts, respectively (**Fig. 4b**). Increased variations in the measured *D* values were also noted in the presence of actin (**Fig. 4b**), likely related to the observed spatial inhomogeneity in local actin density (**Fig. 4a**). These results underscore the key role of actin cytoskeleton in the size-dependent suppression of diffusion in the cytoplasm. We also compared the positively charged HEWL, and noted that whereas its *D* value in the actin-inhibited extract already started low at 14% of the PBS value, a further suppression to 9% of the PBS value was observed in the actin-intact extract (**Fig. S6**).

## Conclusions

Diffusion properties within the complex intracellular environment have been difficult to elucidate, with nuances in molecular behaviors easily masked by subcellular and cell-to-cell heterogeneities. *Xenopus* egg extracts offer a homogenized, near-native cytoplasm system that can be closely examined and readily manipulated. By enabling high-throughput single-molecule statistics, SM*d*M quantifies diffusivity with ∼1% precision at sub-nM diffuser concentrations. Integration of SM*d*M with the extract system thus provides a powerful path toward understanding molecular interactions inside the cell.

Starting with 15 proteins of diverse properties and origins, we observed that whereas the in-extract diffusivities of the negatively charged species displayed a clear size trend at ∼50% of their in-PBS values, the positively charged species diffused substantially more slowly. While the former observation indicates that the diffusion of negatively charged proteins is scaled by a ∼2x increase in cytoplasm viscosity relative to water (which we separately confirmed through bulk viscosity measurements), the latter suggests further diffusion suppression driven by charge-based interactions. Notably, increasing the sample ionic strength effectively alleviated, but did not fully eliminate, this charge-based suppression, indicating that electrostatic interactions between macromolecules are only partially screened even at relatively high salt concentrations. Examination of the mass spectrometry-ranked major proteins in the extract identified most as negatively charged. Although ribosomal proteins tend to be positively charged, their assembly with rRNA results in negatively charged ribosomes. A predominately negatively charged macromolecule environment thus likely drives the observed biased diffusion suppression for positively charged proteins, with the specific degrees of suppression likely dictated by multiple factors, including molecular size, structure, and the amount and distribution of net charges.

Incubation of the extract with RNases could be used to distinguish the contribution of ribosomal proteins and RNA. Interestingly, while RNA degradation led to a moderate increase in diffusivity, a more dramatic effect was the gradual formation of aggregates, likely due to the breakdown of ribosomes and release of >1 mg/mL positively charged ribosomal proteins in the extract, which we reasoned could induce aggregation when surrounded by the abundant negatively charged macromolecules. Adding 1 mg/mL positively charged proteins to the extract induced immediate aggregation, whereas RNase treatments of the ribosome-depleted HSE did not lead to aggregation, further indicating the importance of maintaining a negatively charged macromolecular environment in the cytoplasm, and calling for reexamination of previous interpretations of results involving RNase treatments.

In addition, our experiments underscore the role of the actin cytoskeleton in mediating size-dependent suppression of protein diffusion. Altogether, our results shed new light on how differently charged and sized proteins interact with the complex cytoplasmic environment containing predominantly negatively charged macromolecules and crosslinked meshworks. Besides providing fundamental insights into the inner workings of the cell, the results and methodologies reported in this work may prove valuable in designing *in vitro* systems to dissect the role of cytoplasmic properties and molecular diffusion in diverse cellular processes.

## Materials and Methods

### Xenopus egg cytoplasm extracts

*Xenopus* egg cytoplasm extracts were prepared following standard protocols (*17*). Briefly, eggs from *Xenopus laevis* were dejellied, packed at low speed to remove excess buffer, and then crushed by centrifugation at 18,000 g for 15 min. The cytoplasm was retrieved using a syringe and supplemented with 10 μg/mL LPC protease inhibitors, 20 μg/mL cytochalasin B, and 1x energy mix (4 mM creatine phosphate, 0.5 mM ATP, 0.05 mM EGTA, and 0.5 mM MgCl_2_). Actin-intact extracts were similarly prepared but excluded cytochalasin B. Actin-supplemented extracts were prepared by adding actin from rabbit skeletal muscle (Cytoskeleton, Inc. AKL95-B) to the actin-intact extract to 5 mg/mL. For RNase treatment, Ribonuclease A from bovine pancreas (Sigma R5500) was added to the extract to 1 mg/mL. For the addition of different proteins, poly(D,L-lysine hydrobromide)_250_ (Alamanda Polymers 000-RKB250), HEWL (Sigma L4919), poly(D,L-glutamic acid sodium salt)_300_ (Alamanda Polymers 000-RE300), or BSA (Sigma A3059) were separately added to the extract to 1 mg/mL. High-speed extracts (HSEs) were prepared by ultracentrifuging the standard extracts above at 200,000*g* for 2.5 h at 4 °C. After centrifugation, the clear supernatant was retrieved using a syringe, leaving the pellet behind.

### Dye-labeling of proteins

Sources of proteins used are shown in **Table S1**. We chose proteins of well-defined sizes, many of which have been employed as protein size standards in size-exclusion chromatography (*e*.*g*., Sigma MWGF1000). The protein sizes were further validated with a different set of size standards (Bio-Rad 1511901) (Fig. S7). The proteins were labeled with Cy3B NHS (*N*-hydroxysuccimidyl) ester (Cytiva PA63101) or CF647 NHS ester (Biotium 92135) in 0.1 M NaHCO_3_ at an ∼1:1 initial dye:protein ratio for 1 h. Unconjugated dye was removed by filtering through Amicon centrifugal filters six times, so that no remaining dyes were detected in the final flowthrough. Absorbances at 280 and 560 nm, as measured by a NanoDrop 2000c spectrometer (ThermoFisher), indicated that the final product had ∼0.5 dyes per protein on average. Thus, the fluorescently detected molecules in SM*d*M typically had only one dye on the protein, while the unlabeled fraction of the protein was undetected. Succinylation of Cy3B-labeled HEWL was done by adding excessive solid succinic anhydride (Fisher Scientific AC158760050) into the sample (*57*) and then cleaning up with Amicon centrifugal filters.

### Imaging devices

#1.5 Glass coverslips were acid-treated and passivated with 10 mg/mL methoxy-PEG silane (MW 5000, PG1-SL-5k, Nanocs) in 95% ethanol/water for 30 min, and then rinsed and sonicated for 5 min in Milli-Q water (*30*). The coverslips were then each mounted with a plastic tube (cut from a 0.65 mL microcentrifuge tube) to form an imaging chamber (*58*). For SM*d*M, ∼100 μL of extract or PBS, with dye-labeled proteins added at 0.2-1.0 nM, were added into the imaging chamber. We separately compared imaging chambers with Bioinert surfaces (ibidi 80800), which have been used in recent studies (*27*), and obtained indistinguishable results.

**SM*d*M**. SM*d*M was performed on a Nikon Ti-E inverted fluorescence microscope, as described previously (*8, 30*). Briefly, a 561 nm and a 642 nm laser were focused at the edge of the back focal plane of an oil-immersion objective lens to illuminate a few micrometers into the sample. The focal plane was maintained at ∼3 μm into the sample, and single-molecule images were recorded in the wide field with an EM-CCD camera (iXon Ultra 897, Andor) that operated continuously at 110 frames per second. The laser was repeatedly modulated as paired stroboscopic pulses across tandem camera frames at a fixed center-to-center separation of Δ*t* = 1 ms (**Fig. 1b**). The recorded images across the pair frames thus captured the transient single-molecule displacements in the Δ*t* = 1 ms time window. Typical runs executed 1.0×10^4^ frame pairs over ∼3 min, accumulating 0.5-5×10^5^ single-molecule displacements. These displacements were pooled for global fitting or spatially binned with a grid size of 500 nm for local fitting to the probability model:

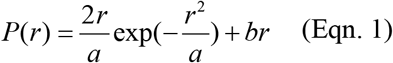

Here *a* = 4*D*Δ*t* with *D* being the diffusion coefficient, and *b* accounts for a uniform background due to extraneous molecules that randomly diffuse into the search radius during Δ*t* (*8*). For spatially binned data, the fitted *D* value in each spatial bin was assigned a color on the continuous *D* scale for rendering into a color-coded *D* map.

### 3D-STORM imaging of actin filaments

For 3D-STORM imaging of actin filaments (*54*) in the actin-intact and actin-added extracts, 100 μL of the extract was added into an 8-well glass coverslip chamber (ibidi 80807) and incubated for 45 min at room temperature. The sample was fixed with 2% glutaraldehyde in the cytoskeleton buffer (10 mM MES, pH = 6.1, 150 mM NaCl, 5 mM EGTA, 5 mM glucose, and 5 mM MgCl_2_) for 30 min, and then reduced using freshly made 0.1% (w/v) NaBH_4_ in PBS. Actin filaments were stained by 400 nM phalloidin Alexa Fluor 647 (Cell Signaling Technology 8940S) overnight at 4 C. The sample was washed with PBS and mounted in a STORM imaging buffer containing 100 mM Tris-HCl, pH = 7.5, 100 mM cysteamine, 5% (w/v) glucose, 0.8 mg/mL glucose oxidase (Sigma-Aldrich, G2133), and 40 μg/mL catalase (Sigma-Aldrich, C30). 3D-STORM was performed as described previously (*55, 56, 59*).

## Acknowledgments

This work was supported by the National Institute of General Medical Sciences of the National Institutes of Health (KX: R35GM149349, RH: R35GM118183), the Packard Fellowships for Science and Engineering (KX), the Heising-Simons Faculty Fellows Award (KX), the Flora Lamson Hewlett Chair in Biochemistry (RH), and a Jane Coffin Childs Memorial Fund Fellowship (CZ).

