## Supplement for "Single-molecule diffusivity quantification in *Xenopus* egg extracts elucidates physicochemical properties of the cytoplasm"

**Table S1.** Diffuser proteins used in this work, their Uniprot IDs, molecular weights (MWs), estimated net charges, and sources. Net charges are estimated using the canonical "Chain" sequences (signal peptides removed) according to Uniprot. Charge estimation is performed with Prot pi (https://www.protpi.ch/) for pH 7.7 (typical pH of the *Xenopus* egg cytoplasm and extract (Good and Heald 2018)) using the ExPASy isoelectric point values, with disulfide bonds applied globally. A shift of −1 net charge is assessed for samples labeled by Cy3B NHS ester. Whereas Cy3B NHS ester is charge-neutral, its conjugation to lysine removes one positive charge on the labeled protein and so shifts the net charge by −1.

| Label in figures | Protein | Uniprot ID | MW (kDa) | Charge @pH7.7 | +Cy3B MW(kDa) | +Cy3B Charge | Provider | Cat# |
|---|---|---|---|---|---|---|---|---|
| **Apoferritin** | Apoferritin, equine (24-mer) | P02791 Q8MIP0 | 480 | -235 | 480 | -236 | Sigma | A3660 |
| **THYG** | Thyroglobulin, bovine (dimer) | P01267 | 660 | -83 | 660 | -84 | Sigma | T1001 |
| **BMY1** | Beta amylase, sweet potato (tetramer) | P10537 | 224 | -59 | 224 | -60 | Sigma | A8781 |
| **ADH** | Alcohol dehydrogenase, yeast (tetramer) | P00330 | 150 | -19.4 | 151 | -20.4 | Sigma | A8656 |
| **BSA** | Bovine serum albumin | P02769 | 66.4 | -17.7 | 67 | -18.7 | Sigma | A3059 |
| **STI** | Soybean trypsin inhibitor | P01070 | 20.0 | -11.6 | 20.6 | -12.6 | Sigma | T2327 |
| **BCA** | Bovine carbonic anhydrase | P00921 | 29.1 | -3.8 | 29.7 | -4.8 | Sigma | C2624 |
| **sHEWL*** | Succinylated HEWL | - | 14.8 | -2.6 | 15.4 | -3.6 | From HEWL | From HEWL |
| **CTRA** | Chymotrypsinogen A, bovine | P00766 | 25.7 | 3.3 | 26.3 | 2.3 | Sigma | C4879 |
| **RNase A** | RNase A, bovine | P61823 | 13.7 | 3.4 | 14.3 | 2.4 | Sigma | R5500 |
| **HEWL** | Lysozyme from chicken egg white | P00698 | 14.3 | 7.4 | 14.9 | 6.4 | Sigma | L4919 |
| **Avidin** | Avidin from chicken egg white (tetramer) | P02701 | 68 | 21.6 | 68.6 | 20.6 | Sigma | A9275 |
| **Cas9-NLS**[#] | Cas9, recombinant, with 1X NLS | Q99ZW2 | 160 | 24.0 | 161 | 23.0 | Sigma | CAS9PROT |
| **H1.0** | Histone H1.0, human, recombinant | P07305 | 20.7 | 52 | 21.3 | 51 | NEB | M2501S |
| **PolyK** | poly(D,L-lysine hydrobromide)$_{250}$ | - | 52 | 250 | 52.6 | 250 | Alamanda Polymers | 000-RKB250 |

*sHEWL: Assuming Cy3B NHS ester conjugated to one of the 6 lysine residues, while succinylation of the remaining 5 lysine residues inversed five +1 charges to −1 (Gitlin *et al.* 2006). This still leaves one N-terminus amine that may be succinylated.

[#]Cas9-NLS: Based on the sequence of Cas9 plus the +5 charged typical nuclear localization sequence (NLS) PKKKRKV.

**Table S2.** List of the most abundant cytoplasmic proteins in the *Xenopus* egg cytoplasmic extract and their estimated net charge $q$. Cytoplasmic proteins (not including RNA-binding and ribosomal proteins in Table S3) are ordered by their estimated concentrations $c$ in the extract according to the mass spectrometry data of (Wühr *et al.* 2014), based on updated annotations by Wühr Lab (https://wuehr.scholar.princeton.edu/protein-concentrations-xenopus-egg). Net charges at pH 7.7 are estimated for the annotated sequences as described in Table S1. Most proteins on this list are negatively charged or neutral. A few proteins in the glycolysis pathway (GAPDH and AldOa) are weakly positively charged (+1 to +3), which may mediate binding to negatively charged phosphorylated glucose metabolites, thereby neutralizing the total charge.

| ***Laevis* gene** | **Human gene** | **Protein description** | $c$ **(μM)** | **Net $q$** |
|---|---|---|---|---|
| serpina6.L | SERPINA1 | Alpha-1-antitrypsin | 8.7 | 0.2 |
| actc1.L | ACTC1 | Actin, alpha cardiac muscle 1 | 8.0 | -13.8 |
| eno1.L | ENO1 | Alpha-enolase | 7.9 | -6.9 |
| cgl.2.L | ITLN2 | Intelectin-2 | 6.3 | -7.7 |
| hbz.L | HBZ | Hemoglobin subunit zeta | 5.9 | -4.8 |
| actg1.S | ACTG1 | Actin, cytoplasmic 2 | 5.8 | -12.8 |
| gapdh.L | GAPDH | Glyceraldehyde-3-phosphate dehydrogenase | 5.8 | 2.1 |
| tpi1.S | TPI1 | Triosephosphate isomerase | 5.8 | -0.9 |
| ldhb.S | LDHB | L-lactate dehydrogenase B chain | 5.5 | -6.6 |
| nme2.S | NME2 | Nucleoside diphosphate kinase B | 5.3 | -1.9 |
| fabp4 | FABP4 | Fatty acid-binding protein, adipocyte | 4.6 | -1.9 |
| rps27a.S* | RPS27A | Ubiquitin | 4.4 | -0.9 |
| pkm.L | PKM | Pyruvate kinase PKM | 4.2 | -8.8 |
| plin2.L | PLIN2 | Perilipin-2 | 4.2 | -16.9 |
| grhpr.2.L | GRHPR | Glyoxylate reductase/hydroxypyruvate reductase | 4.1 | -4.9 |
| nme2.L | NME2 | Nucleoside diphosphate kinase B | 4.1 | -1.9 |
| mdh1.S | MDH1 | Malate dehydrogenase, cytoplasmic | 4.1 | -1.9 |
| hba-l5.L | HBZ | Hemoglobin subunit zeta | 4.0 | -4.8 |
| ckb.S | CKB | Creatine kinase B-type | 3.9 | -7.8 |
| pgk1.L | PGK1 | Phosphoglycerate kinase 1 | 3.9 | -1.9 |
| gapdh.S | GAPDH | Glyceraldehyde-3-phosphate dehydrogenase | 3.9 | 1.1 |
| aldoa.S | ALDOA | Fructose-bisphosphate aldolase A | 3.7 | 3.2 |
| gstm1.S | GSTM1 | Glutathione S-transferase Mu 1 | 3.7 | 0.1 |
| hpgds.S | HPGDS | Hematopoietic prostaglandin D synthase | 3.6 | -2.0 |
| ahcy.S | AHCY | Adenosylhomocysteinase | 3.6 | -6.9 |
| ftmt.S | FTH1 | Ferritin heavy chain | 3.5 | -11.7 |
| gpi.L | GPI | Glucose-6-phosphate isomerase | 3.4 | -2.7 |
| tpi1.L | TPI1 | Triosephosphate isomerase | 3.3 | -1.9 |

* rps27a.S (RPS27A) encodes both ubiquitin and the 40S ribosomal protein S27a. The charge here is calculated for ubiquitin.

**Table S3.** List of the most abundant RNA-binding and ribosomal proteins in the *Xenopus* egg cytoplasmic extract as determined by mass spectrometry (Wühr *et al.* 2014), and their estimated net charges, generated as in Table S2. Note that when assembled with rRNA into ribosomes, the strong negative charges on rRNA (−1 net charge per nucleotide on the RNA backbone) overcompensate for protein charges to render the assembled ribosomes highly negatively charged (Knight *et al.* 2013; Schavemaker *et al.* 2017).

| ***Laevis* gene** | **Human gene** | **Protein description** | ***c* (µM)** | **Net *q*** |
|---|---|---|---|---|
| rpl32.L | RPL32 | 60S ribosomal protein L32 | 5.0 | 25.1 |
| rpl23a.S | RPL23A | 60S ribosomal protein L23a (Fragment) | 4.7 | 25.1 |
| rps27a.S* | RPS27A | 40S ribosomal protein S27a | 4.4 | 15.4 |
| cirbp.S | CIRBP | Cold-inducible RNA-binding protein | 4.1 | 2.1 |
| rps14.L | RPS14 | 40S ribosomal protein S14 | 4.1 | 12.2 |
| rps9 | RPS9 | 40S ribosomal protein S9 | 4.0 | 21.1 |
| rpl27.S | RPL27 | 60S ribosomal protein L27 | 3.9 | 24.0 |
| rpl7a.S | RPL7A | 60S ribosomal protein L7a | 3.8 | 40.0 |
| rps18 | RPS18 | 40S ribosomal protein S18 | 3.8 | 33.1 |
| rps16.S | RPS16 | 40S ribosomal protein S16 | 3.8 | 16.1 |
| rps3.L | RPS3 | 40S ribosomal protein S3 | 3.7 | 11.1 |
| rps8.L | RPS8 | 40S ribosomal protein S8 | 3.7 | 33.1 |
| rps28p9.L | RPS28 | 40S ribosomal protein S28 | 3.6 | 4.2 |
| LOC398653 | RPLP2 | 60S acidic ribosomal protein P2 | 3.5 | -8.9 |
| rplp0.S | RPLP0 | 60S acidic ribosomal protein P0 | 3.5 | -6.9 |
| rps4x.L | RPS4X | 40S ribosomal protein S4, X isoform | 3.4 | 23.2 |
| rps12.L | RPS12 | 40S ribosomal protein S12 | 3.4 | -0.9 |
| rpl27a.S | RPL27A | 60S ribosomal protein L27a | 3.4 | 22.3 |
| rpl26.L | RPL26 | 60S ribosomal protein L26 | 3.3 | 24.1 |
| rpl22.L | RPL22 | 60S ribosomal protein L22 | 3.3 | 6.1 |
| rpl8.L | RPL8 | 60S ribosomal protein L8 | 3.3 | 36.2 |
| rps19.S | RPS19 | 40S ribosomal protein S19 | 3.3 | 16.2 |
| rps17.S | RPS17 | 40S ribosomal protein S17 | 3.3 | 8.1 |
| rps25.L | RPS25 | 40S ribosomal protein S25 | 3.3 | 20.0 |
| rps11 | RPS11 | 40S ribosomal protein S11 | 3.1 | 22.1 |
| rpl19 | RPL19 | 60S ribosomal protein L19 | 3.1 | 43.1 |
| rpl7.L | RPL7 | 60S ribosomal protein L7 | 3.1 | 41.0 |
| rps6.L | RPS6 | 40S ribosomal protein S6 | 3.0 | 40.0 |
| rpl4.S | RPL4 | 60S ribosomal protein L4 | 3.0 | 63.0 |

* rps27a.S (RPS27A) encodes both ubiquitin and the 40S ribosomal protein S27a. The charge here is calculated for the 40S ribosomal protein S27a.

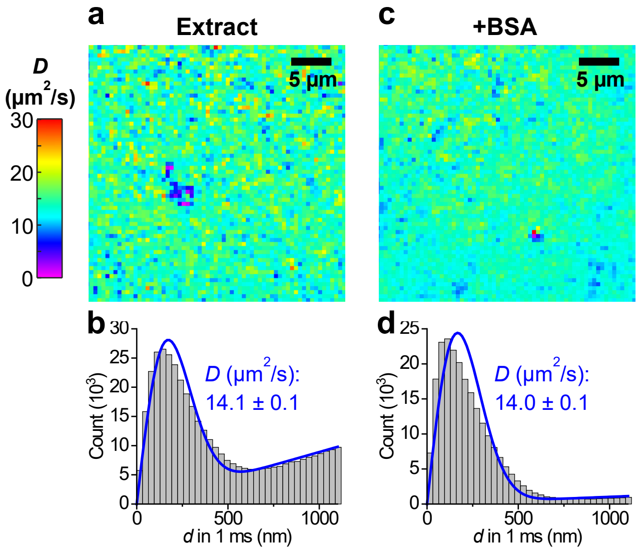


**Figure S1.** SM*d*M diffusivity mapping of Cy3B-labeled HEWL in untreated and BSA-supplemented *Xenopus* egg extracts. (**a**) Color-coded SM*d*M diffusivity map for an untreated sample, presented on the same color scale as **Fig. 3**. (**b**) Distribution of the 1-ms single-molecule displacements from (a). Blue curve: fit to our single-mode SM*d*M diffusion model, with resultant apparent diffusion coefficients *D* and 95% confidence intervals marked. Note that in this sample, a higher concentration of Cy3B-labeled HEWL (~1 nM) was used to increase the detected single-molecule density to facilitate spatial mapping. While this led to a higher background in the displacement distribution (when compared to **Fig. 1d**), the *D* value obtained with our fitting model (Eqn. 1) stays consistent, in agreement with what we have analyzed and demonstrated previously (Xiang *et al.* 2020). (**c**) Color-coded SM*d*M diffusivity map for another sample to which BSA was added at 1 mg/mL. (**d**) Distribution of the 1-ms single-molecule displacements from (c) and fit to our model.

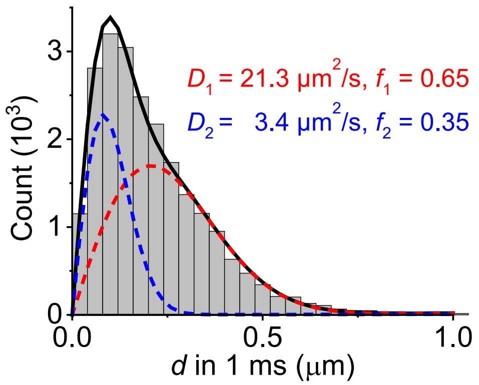


**Figure S2.** Two-component fit to the distribution of SM*d*M-recorded 1-ms single-molecule displacements of Cy3B-labeled HEWL diffusing in extract as shown in Fig. 1d. Histogram: displacements; Red curve: fast component of the fit ($D_1$ = 21.3 μm$^2$/s, fraction $f_1$ = 0.65); Blue curve: slow component of the fit ($D_2$ = 3.4 μm$^2$/s, fraction $f_2$ = 0.35); Black curve: sum of the red and blue curves. While the two-component fit improves over the single-component model (Fig. 1d), the system may be better assumed as a continuous distribution of different transient states.

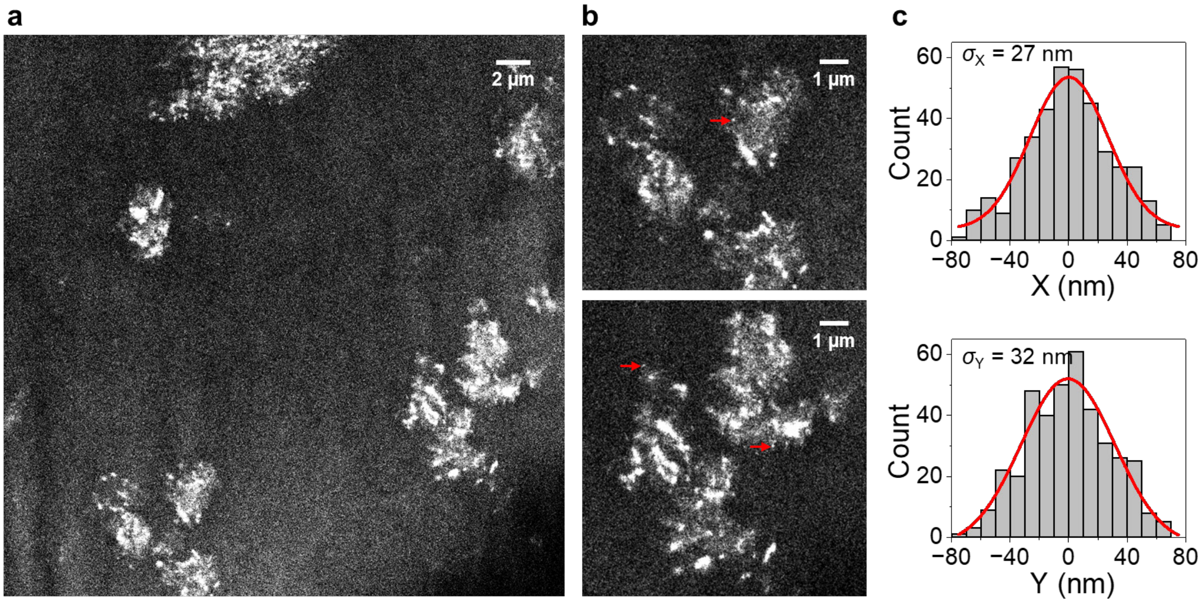


**Figure S3.** SMLM super-resolution images of Cy3B-labeled HEWL in RNase-treated extract, as generated from the single-molecule localizations of the SM*d*M data. (a) Same field of view as the SM*d*M data of Fig. 3c. (b) Zoom-ins of two regions. (c) Distribution of single-molecule positions for overlaid 4 small nanoclusters like those indicated by the red arrows in (b), in the X (top) and Y (bottom) directions, respectively. Gaussian fits (red curves) give standard deviations of 27 and 32 nm.

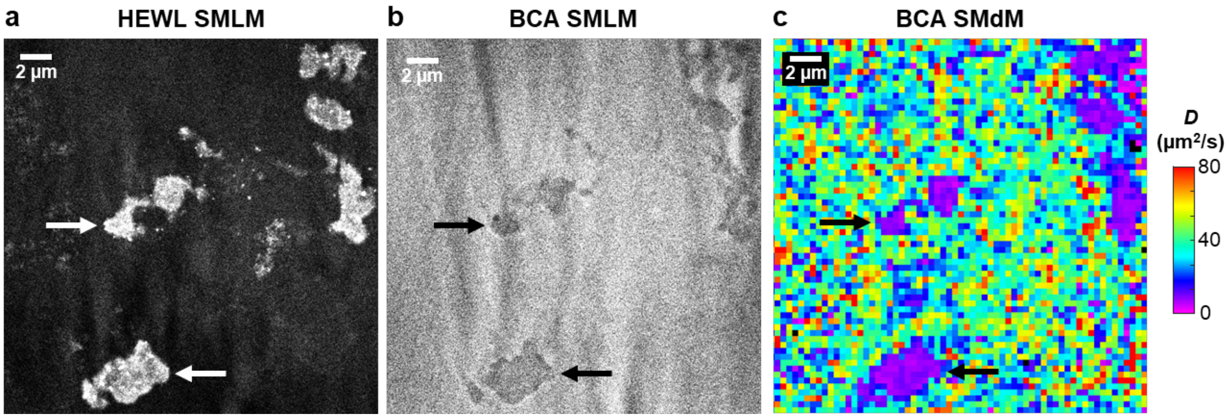


**Figure S4.** Diffusion of the negatively charged BCA in RNase-treated extract. (a,b) SMLM images of Cy3B-labeled HEWL (a) and CF647-labeled BCA (b) in an RNase-treated extract sample. Vertical stripe patterns are attributed to local lensing effects from the high refractive indices of the aggregates under our inclined illumination scheme (He *et al.* 2023). (c) SM*d*M diffusivity map of the CF647-labeled BCA. Arrows point to aggregates, where diffusivity reduction was accompanied by local increases and decreases in the abundances of HEWL and BCA, respectively. These results may be interpreted as that as the RNase-released positively charged proteins interact with the negatively charged cytoplasm environment, the resultant aggregates are overwhelmed by the latter on the surface. Positively and natively charged tracer proteins are thus respectively attracted to and repelled by the aggregates, with the former further likely participating in the aggregate cores.

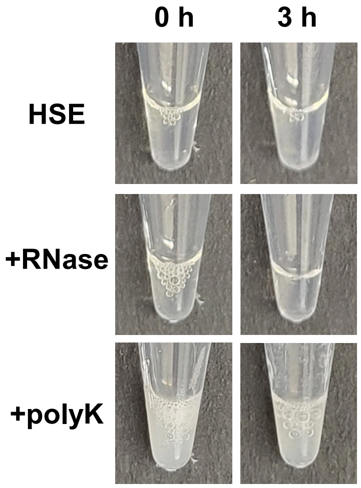


**Figure S5.** Aggregation assays for the ribosome-depleted high-speed extracts (HSEs). Photos are shown for HSEs at 0 h (left) and 3 h (right), without or with the addition of RNase or 1 mg/mL polylysine. Whereas polylysine addition generated immediate clouding, RNase treatment did not induce clouding over 3 h, consistent with our model that in the crude, ribosome-containing cytoplasm extract, RNase degradation of rRNA releases positively charged ribosomal proteins to cause aggregation.

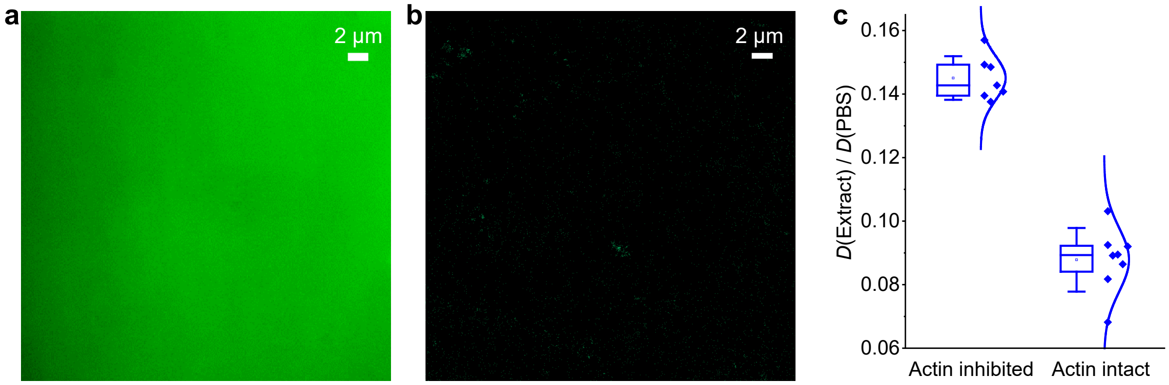


**Figure S6.** Additional data related to actin filaments. (a,c) Typical epifluorescence (a) and STORM (b) images of phalloidin-AF647 in actin-inhibited *Xenopus* egg extracts, showing no resolvable structures. (c) SM*d*M-determined *D* values in the extract relative to in PBS, for the positively charged HEWL in actin-inhibited and actin-intact extracts. Each data point corresponds to one independent SM*d*M measurement for a different sample region, from 5 actin-inhibited and 3 actin-intact samples, respectively.

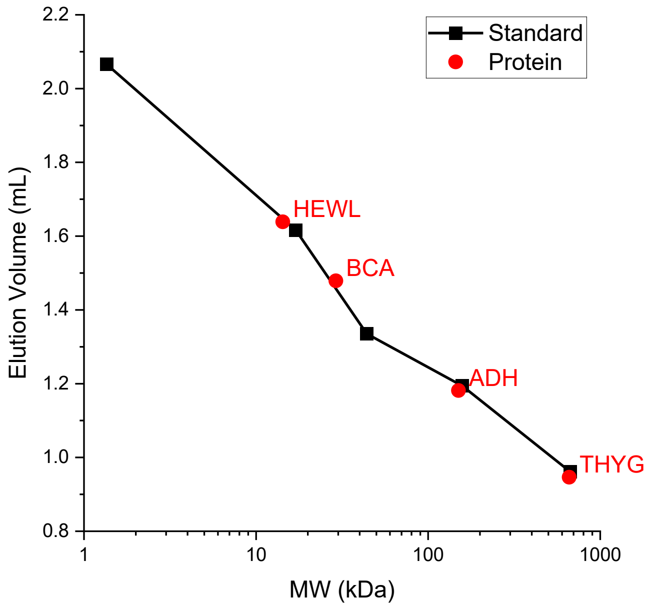


**Figure S7.** Size-exclusion chromatography elution volumes of four of the protein samples used in this study (red circles) compared to calibration standards (Bio-Rad 1511901; black squares), plotted versus the expected molecular weight. Separation was performed at 4 °C in sterile-filtered PBS. 50 µL of protein solution (1 mg/mL) was injected into an ÄKTA pure micro chromatography system (Cytiva 29302479) onto a Superdex 200 Increase 3.2/300 column (Cytiva 28990946) at a flow rate of 0.025 mL/min. The sample elution was monitored by absorbance at 280 nm.
